# The IL-33/ST2 signaling axis drives pathogenesis in acute SARS-CoV-2 infection

**DOI:** 10.1101/2024.11.27.625579

**Authors:** Claire Fleming, Henry J McSorley, Judith E Allen, William A Petri

**Author notes:** Corresponding author: William A. Petri Jr. University of Virginia, 345 Crispell Drive, Charlottesville Virginia 22908-1340, USA.

## Abstract

Severe acute respiratory syndrome coronavirus 2 (SARS-CoV-2), the causative agent of coronavirus disease 2019 (COVID-19), remains a significant threat to global public health. Immunopathological damage plays a role in driving pneumonia, acute respiratory distress syndrome (ARDS), and multiorgan failure in severe COVID-19. Therefore, dissecting the pulmonary immune response to SARS-CoV-2 infection is critical to understand disease pathogenesis and identify immune pathways targetable by therapeutic intervention. Considering that the type 2 cytokine IL-13 enhances COVID-19 disease severity, therapeutic targeting of upstream signals that drive type 2 immunity may confer further protection. In this study, we investigate the role of the IL-33/ST2 signaling axis, a potent inducer of type 2 immunity in the lung, in a mouse model of COVID-19. Upon infection with mouse-adapted SARS-CoV-2 MA10, ST2^-/-^ mice had significantly improved weight loss and survival (69.2% vs 13.3% survival; P = 0.0005), as compared to wild-type mice. In a complementary pharmacologic approach, IL-33/ST2 signaling was inhibited using HpBARI_Hom2, a helminth derived protein that binds to mouse ST2 and blocks IL-33 signaling. In SARS-CoV-2 MA10 infection, HpBARI_Hom2-treated mice had significantly improved weight loss and survival (60% vs 10% survival; P = 0.0035), as compared to inert control-treated mice. These data demonstrate that loss of IL-33/ST2 signaling confers protection during acute SARS-CoV-2 MA10 infection, implicating the IL-33/ST2 signaling axis as an enhancer of COVID-19 disease severity. The protection conferred by pharmacologic blockade of IL-33/ST2 signaling was independent of viral control, as HpBARI_Hom2-treated mice had no reduction in viral titers. This finding suggests an immunopathogenic role for IL-33/ST2 signaling. One potential mechanism through which IL-33/ST2 signaling may drive severe disease is through enhancement of type 2 immune pathways including IL-5 production, as pulmonary IL-5 concentrations were found to depend on IL-33/ST2 signaling in acute SARS-CoV-2 MA10 infection.

## INTRODUCTION

Severe acute respiratory syndrome coronavirus 2 (SARS-CoV-2), the causative agent of coronavirus disease 2019 (COVID-19), has brought the threat of emerging infectious pathogens to the forefront of public attention. Since late 2019, SARS-CoV-2 infection has caused greater than 7 million deaths worldwide^1^. Severe COVID-19 is characterized by systemic hyperinflammation and elevated levels of circulating proinflammatory cytokines^2^. Immunopathological damage drives pneumonia, acute respiratory distress syndrome (ARDS), and multiorgan failure in severe COVID-19^3^. Therefore, dissecting the pulmonary immune response to SARS-CoV-2 infection is critical to understand disease pathogenesis and identify immune pathways targetable by therapeutic intervention. Immunomodulators including broadly anti-inflammatory corticosteroids and more targeted treatments including an interleukin-6 (IL-6) inhibitor (tocilizumab) and a JAK1/2 inhibitor (baricitinib) have shown success in treatment of severe COVID-19^4^. However, in-hospital mortality remains high for patients with severe COVID-19^5^, highlighting the need for continued development of therapeutics. Further, with the emergence of three highly pathogenic coronaviruses within two decades (SARS-CoV, Middle East respiratory syndrome coronavirus (MERS-CoV), and SARS-CoV-2), there is a clear need to develop an arsenal of effective therapeutics to help us prepare for future coronavirus outbreaks.

Targeting of type 2 immune pathways is a potential avenue of therapeutic intervention in severe COVID-19. In particular, research has demonstrated a link between the type 2 cytokine IL-13 and COVID-19 severity^6^. Implicated in several allergic diseases, IL-13 is a potent inducer of type 2 immune pathways including mucus secretion, goblet cell metaplasia, smooth muscle cell contraction, and subepithelial fibrosis^7^. IL-13 was found to be elevated in the plasma of patients with severe COVID-19 and high IL-13 levels were associated with an increased probability of requiring mechanical ventilation^8^. In addition to serving as a biomarker for severe disease, therapeutic targeting of IL-13 signaling has shown promise in both mouse models and human patients. IL-13 neutralization was found to be protective in the K18-hACE2 mouse model of COVID-19^8^ and in a phase 2a clinical trial, treatment with dupilumab (an antibody that blocks IL-4Rα, the shared receptor for IL-4 and IL-13) increased survival^9^ and improved pulmonary function tests at one year follow-up^10^ (secondary outcomes). What remains to be determined is what signals are responsible for the induction of type 2 immunity in the context of this respiratory viral infection and whether therapeutic intervention targeting such upstream signals would confer further therapeutic benefit.

One signal potentially responsible for the induction of type 2 immunity during acute SARS-CoV-2 infection is the alarmin cytokine IL-33. A member of the IL-1 family of cytokines, IL-33 is stored in the nucleus of epithelial, endothelial, and stromal cells under steady-state conditions^11^. Upon cellular damage or tissue injury, IL-33 is released into the extracellular space^11^. IL-33 binds to a heterodimeric receptor composed of serum stimulated gene 2 (ST2) and IL-1 receptor accessory protein (IL-1RAcP). While the IL-1RAcP co-receptor is shared among other IL-1 family members, ST2 binds specifically to IL-33^12^. IL-33 is a potent inducer of type 2 immune pathways in the lung, as ST2 is constitutively expressed by tissue-resident type 2 cells including mast cells and type 2 innate lymphoid cells (ILC2s)^13^. Considered to be the innate counterpart to T helper (Th) cells, ILCs play a major role in the early immune response through secretion of effector cytokines and ILC2s are the primary innate source of IL-13 in the lungs^14^. Other IL-33 targets include eosinophils, basophils, neutrophils, macrophages, dendritic cells (DCs), Th1 cells, Th2 cells, CD8+ T cells, regulatory T cells (Tregs), and natural killer (NK) cells; while several of these cell types express ST2 constitutively, others selectively express ST2 when exposed to an inflammatory microenviornment^11^.

In COVID-19 patients, elevated levels of circulating IL-33 are known to be predictive of poor outcomes^15^ or associated with severe disease^16,17^. Though it failed to meet its primary endpoint (time to clinical response), a recent phase 2a trial evaluating the IL-33 neutralizing antibody tozorakimab in COVID-19 patients showed enhanced efficacy in the subgroup of patients with higher baseline levels of serum IL-33^18^. It has recently been shown in a mouse model of COVID-19 that *IL-33*^*-/-*^ mice experienced reduced weight loss following infection with a mouse adapted SARS-CoV-2 virus (CMA3p20) and this protection correlated with a reduction in innate immune cell infiltrates in the acutely infected lungs^19^. These findings suggest that IL-33 may play an immunopathogenic role in COVID-19, but the mechanism underlying IL-33-mediated pathogenesis remains to be fully understood.

In this study, we investigate the role of IL-33/ST2 signaling in a mouse model of COVID-19. Specifically, we used the SARS-CoV-2 MA10 virus, which was generated using reverse genetic engineering of the viral S protein, allowing the virus to efficiently bind mouse ACE2^20^, followed by subsequent serial *in vivo* passaging in mice^21^. This model shows a dose- and age-related increase in pathogenesis and recapitulates pathological features of acute lung injury (ALI) and ARDS^21^. Using complementary genetic and pharmacologic approaches, we found that the IL-33/ST2 signaling axis enhances disease severity of COVID-19. Interestingly, the protection conferred by pharmacologic blockade of IL-33/ST2 signaling was independent of viral control. This finding suggests an immunopathogenic role for IL-33/ST2 signaling, potentially through enhancement of type 2 immune pathways including IL-5 production.

## RESULTS

To test whether IL-33/ST2 signaling enhances disease severity of COVID-19, we used complementary genetic knockout and pharmacologic approaches to interrupt the IL-33/ST2 signaling axis in a mouse model of COVID-19. First, we infected ST2^-/-^ and wild-type C57BL/6 control mice with SARS-CoV-2 MA10 and measured their survival and weight loss for the subsequent 14 days post-infection. The ST2^-/-^ mice had significantly improved survival (69.2% vs 13.3% survival; P = 0.0005) and weight loss, as compared to wild-type controls (Figure 1A-C). As a complementary pharmacologic approach, we inhibited IL-33/ST2 signaling using a protein named *Heligmosomoides polygyrus* binds alarmin receptor and inhibits homologue 2 (HpBARI_Hom2). This immunomodulatory protein, originally identified from the murine parasite *H. polygyrus*, binds the ST2 receptor with high affinity and blocks the interaction of IL-33 with ST2^22^. We administered HpBARI_Hom2 to C57BL/6 mice via intranasal administration on the day prior to SARS-CoV-2 MA10 infection, with follow-up intraperitoneal administration on the day of infection and days 2 and 5 post-infection. HpBARI_Hom2-treated mice had improved survival (60% vs 10% survival; P = 0.0035) and weight loss, as compared to mice that were treated with an inert control protein (enzymatically inactive *H. polygyrus* Acetylcholinesterase (HpAChE)) generated in the same recombinant expression system (Figure 1D-F).

**Figure 1:**
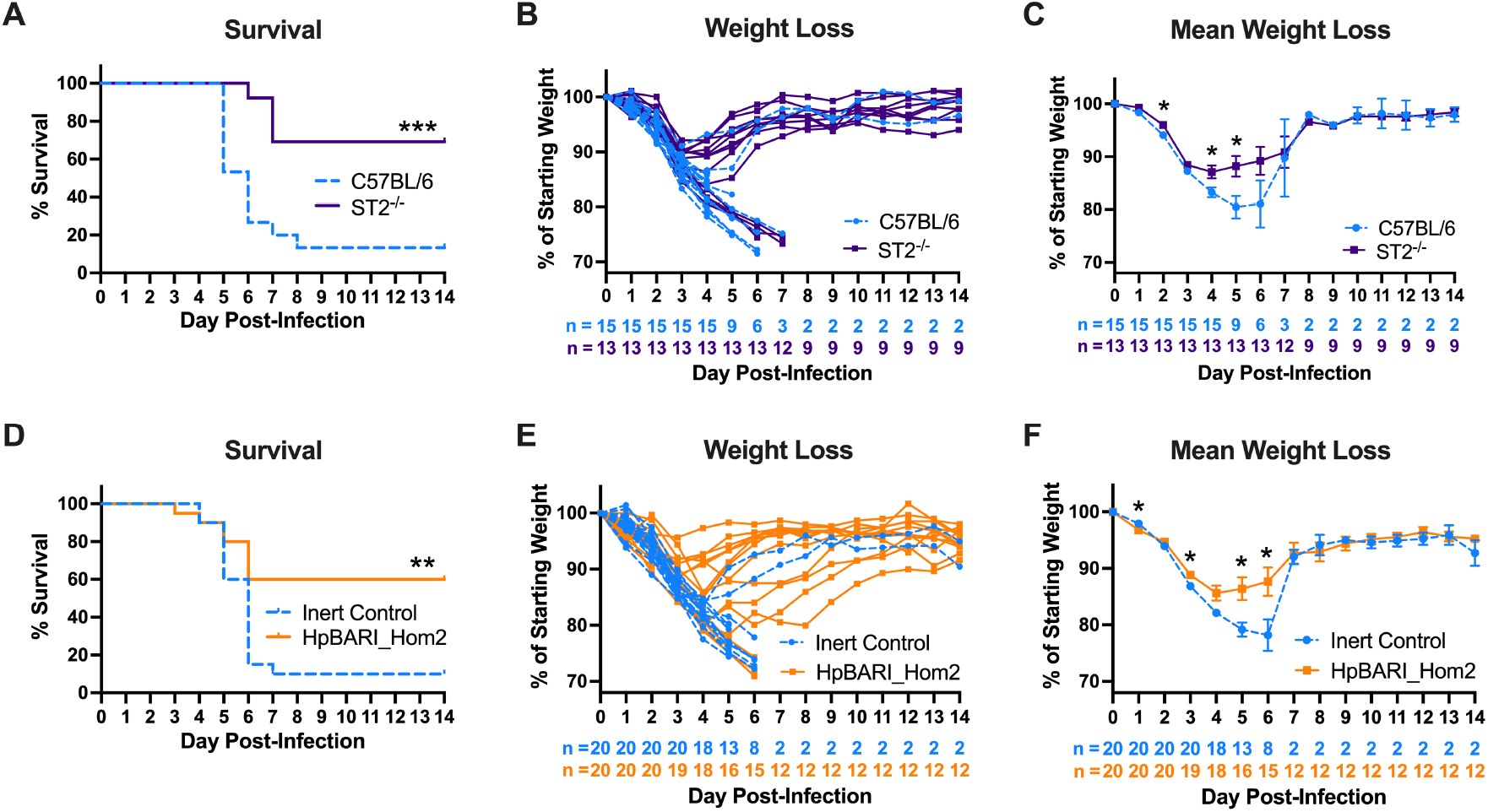
Loss of IL-33/ST2 signaling through genetic knockout or pharmacologic blockade of ST2 improves survival and weight loss in SARS-CoV-2 MA10 infection. C57BL/6 mice were infected with SARS-CoV-2 MA10 on day 0. Kaplan-Meier survival curve (A), weight loss (B), and mean weight loss (C) of ST2^-/-^ mice (purple) and wild-type mice (blue). Kaplan-Meier survival curve (D), weight loss (E), and mean weight loss (F) of mice treated with 10 μg of either ST2-blocking HpBARI_Hom2 (orange) or control inert protein (blue) on days -1, 0, 2, and 5. Data combined from 2 independent experiments. P-values for survival determined by log-rank test. P-values for weight loss determined by non-parametric test. *P<0.05, **P<0.01, ***P<0.001.

To begin to assess the mechanism through which IL-33/ST2 signaling enhances COVID-19 disease severity, we next assessed lung tissue and bronchoalveolar lavage fluid (BALF) of HpBARI_Hom2-treated and inert control-treated mice at a timepoint of acute SARS-CoV-2 MA10 infection. Specifically, we administered HpBARI_Hom2 to C57BL/6 mice via intranasal administration on the day prior to SARS-CoV-2 MA10 infection, with follow-up intraperitoneal administration on the day of infection and day 2 post-infection. We collected lung tissue and BALF at day 4 post-infection. This is a timepoint at which there are clear signs of disease onset in terms of weight loss, but not yet substantial mortality (Figure 1D-F). Viral titers were measured by plaque assay and despite significant protection in survival studies (Figure 1D), HpBARI_Hom2-treated mice had no reduction in viral titers as compared to inert control-treated mice at this timepoint (Figure 2A). HpBARI_Hom2-treated mice also had no significant reduction in lung damage as compared to inert control-treated mice at this timepoint (Figure 2B), as measured by scoring hematoxylin and eosin (H&E)-stained lung sections according to the American Thoracic Society’s ALI scoring system^23^. Considering a potential immunopathogenic role for IL-33/ST2 signaling, a 32-plex cytokine Luminex panel was performed on BALF to investigate how the IL-33/ST2 signaling axis impacts the cytokine environment of the acutely infected lung. Though the cytokine milieu was highly heterogenous between infected mice, HpBARI_Hom2-treated mice had a significant increase in IL-2 and a significant reduction in IL-5 as compared to inert control-treated mice (Figure 2C).

**Figure 2:**
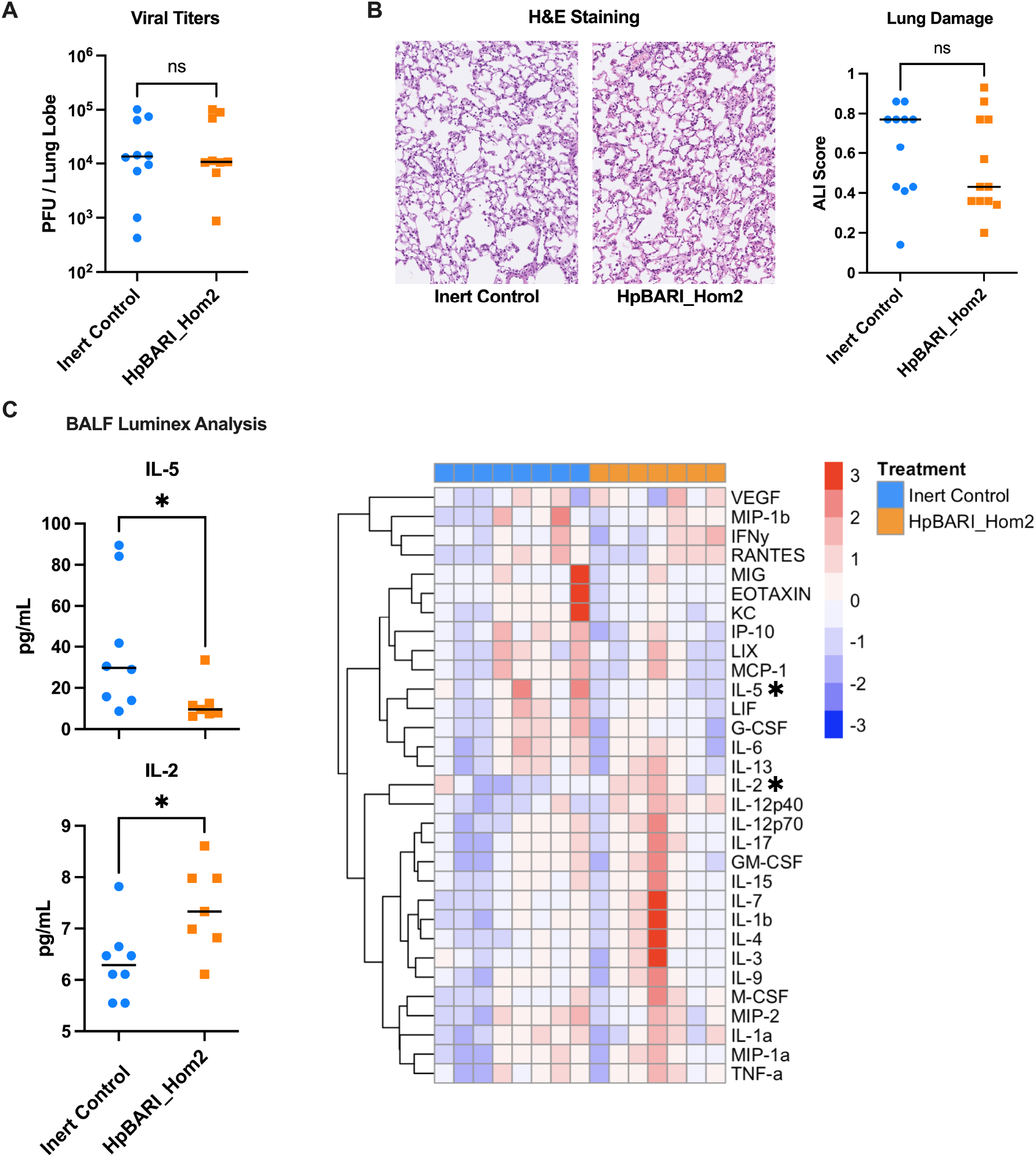
Pharmacologic blockade of ST2 by HpBARI_Hom2 does not impact viral titers or tissue pathology on day 4 post infection but does alter the cytokine environment of the acutely infected lung. C57BL/6 mice were infected with SARS-CoV-2 MA10 on day 0 and were treated with 10 μg of either HpBARI_Hom2 (orange) or control inert protein (blue) on days -1, 0, 2. Mice were sacrificed at day 4 post infection. Lung tissue was collected for measurement of viral titers by plaque assay (A) and evaluation of tissue pathology by scoring H&E slides for ALI (B). BALF was collected and analyzed by a 32-plex Luminex cytokine panel and concentrations are visualized by heatmap, with color scaled by row (C). IL-10 was below the limit of detection for all samples assayed and is excluded from visualization. Viral titer, tissue pathology, and Luminex data combined from 2 independent experiments. P-values determined by non-parametric test. *P<0.05.

## DISCUSSION

In this study, we investigated the role of the IL-33/ST2 signaling axis in a mouse model of COVID-19. Complementary genetic and pharmacologic approaches demonstrated that loss of IL-33/ST2 signaling confers protection during acute SARS-CoV-2 MA10 infection, implicating the IL-33/ST2 signaling axis as an enhancer of COVID-19 disease severity. These findings concur with published work showing *IL-33*^-/-^ mice were protected from severe disease in another mouse model of COVID-19, a non-lethal model using mouse adapted SARS-CoV-2 CMA3p20^19^. ‘While data from COVID-19 patients have shown that elevated levels of circulating IL-33 are predictive of poor outcomes^15^ or associated with severe disease^16,17^, our data, as well as published data from the CMA3p20 mouse model^19^, demonstrate that IL-33 is not solely a useful biomarker, but rather is an important driver of disease pathogenesis.

Though pharmacologic blockade of IL-33/ST2 signaling via HpBARI_Hom2 protected mice from severe COVID-19 disease, HpBARI_Hom2 treatment did not reduce viral titers at a timepoint of acute infection. Interestingly, in studies involving IL-13 neutralization in a mouse model of COVID-19, mice were similarly protected from severe disease without a reduction in viral titers^8^. Collectively, these findings demonstrate that type 2 pathways can drive immunopathogenesis through mechanisms independent of viral control. Considering an immunopathogenic role for IL-33/ST2 signaling, an exploratory Luminex panel was performed to investigate how the IL-33/ST2 signaling axis impacts the cytokine environment of the acutely infected lung. HpBARI_Hom2 treatment was found to increase IL-2 and reduce IL-5 in the BALF at a timepoint of acute infection. The reduction in IL-5 is consistent with the known role of IL-33/ST2 signaling as an enhancer of type 2 immunity. Though interestingly, despite the fact that IL-33/ST2 signaling can induce IL-13 and IL-4 production in several cell types (e.g. ILC2s and basophils), we did not observe any significant difference in these type 2 cytokines with HpBARI_Hom2 treatment at this particular timepoint. It is possible that induction of IL-5 is a mechanism through which IL-33/ST2 signaling drives immunopathogenesis, a hypothesis that will be tested in future mechanistic studies. This work demonstrates IL-33/ST2 signaling enhances disease severity of COVID-19, validating this immune pathway as a potential therapeutic target.

## METHODS

### Virus

All experiments were performed in the University of Virginia biosafety level 3 (BSL-3) laboratory. The SARS-CoV-2 MA10 virus was obtained through “BEI Resources, NIAID, NIH: SARS-Related Coronavirus 2, Mouse-Adapted, MA10 Variant (in isolate USA-WA1/2020 backbone), Infectious Clone (ic2019-nCoV MA10) in Calu-3 Cells, NR-55329, contributed by Ralph S. Baric”. Upon receipt, the virus was propagated in Vero C1008, Clone E6 (ATCC CRL-1586) cells cultured in Dulbecco’s Modified Eagle’s Medium (DMEM) supplemented with 10% fetal bovine serum (FBS) and grown at 37°C, 5% CO_2_. The passage 1 (P1) stocks generated were then used to infect additional Vero E6 cells generating passage 2 (P2) stocks which were used for all experiments.

### Mouse infections

All animal experiments were approved by the Institutional Animal Care and Use Committee (IACUC) at the University of Virginia and were performed in the animal biosafety level 3 (ABSL-3) laboratory. For survival experiments, mice were anesthetized with ketamine/xylazine and infected with 5.5 × 10^3^ PFU of SARS-CoV-2 MA10 by intranasal administration. The mice were evaluated for 14 days post-infection and were monitored daily for weight loss. If during the course of infection mice reached a predetermined reduction in body weight, they were euthanized. For other experiments, mice were infected with 5.5 × 10^3^ - 1.5 × 10^4^ PFU of SARS-CoV-2 MA10 and were monitored daily until euthanized at day 4 post-infection for evaluation of pulmonary tissue by downstream analyses (plaque assay, histology, viral titers, and Luminex analysis).

### Genetic knockout of IL-33/ST2 signaling

For experiments involving genetic knockout of IL-33/ST2 signaling, male C57BL/6 and ST2^-/-^ mice on the C57BL/6 background were used. Mice were age-matched within a week between experimental groups and were infected at 18-21 weeks old. C57BL/6 were purchased from Charles River and were aged for at least a month in the University of Virginia vivarium to equilibrate their microbiota. ST2^-/-^ mice (originally developed as described^24^) were bred in the University of Virginia vivarium.

### Pharmacologic blockade of IL-33/ST2 signaling

For experiments involving pharmacologic blockade of IL-33/ST2 signaling, male C57BL/6 mice were purchased from Jackson Laboratory and were infected at 20-21 weeks-old. IL-33/ST2 signaling was inhibited using a protein named *H. polygyrus* binds alarmin receptor and inhibits homologue 2 (HpBARI_Hom2). This immunomodulatory protein expressed by the murine parasite *H. polygyrus* binds the ST2 receptor with high affinity and blocks the interaction of IL-33 with ST2^22^. For these experiments, HpBARI_Hom2 recombinantly expressed in HEK293 cells and biochemically purified using affinity purification techniques was provided by Dr. Henry McSorley (University of Dundee). To account for potential non-specific immunomodulatory impacts that introduction of a recombinant protein may induce, control mice were treated with an inert protein (enzymatically inactive *H. polygyrus* acetylcholinesterase (HpAchE)) generated in the same recombinant expression system and also provided by Dr. McSorley (University of Dundee). On the day prior to SARS-CoV-2 MA10 infection (day -1), mice were anesthetized with isoflurane and 10 μg of either ST2-blocking HpBARI_Hom2 or the control inert protein was administered intranasally. For survival experiments, three additional doses of 10 μg HpBARI_Hom2 or inert control were given on the day of infection (day 0) and days 2 and 5 post-infection via intraperitoneal injection. For experiments in which mice were euthanized at day 4 post-infection, only two additional doses of 10 μg HpBARI_Hom2 or inert control were given on days 0 and 2.

### Plaque Assays

In these experiments, mice were infected with 5.5 × 10^3^ PFU SARS-CoV-2 MA10 and were euthanized at day 4 post-infection. The right caudal lung lobe was homogenized in 0.5 mL serum-free DMEM with a disposable tissue grinder. The homogenate was centrifuged at 400 ×g for 7 minutes at 4°C and the supernatant was collected and stored at -80°C. To quantify viral load by plaque assay, Vero C1008 Clone E6 (ATCC CRL-1586) cells cultured in DMEM with 10% FBS were seeded in 12-well plates at a concentration of 2 × 10^5^ cells/well. The next day, serial dilutions of the stored lung homogenates were prepared in serum-free DMEM. Following media aspiration, the confluent Vero E6 cells were rinsed with sterile PBS prior to addition of the diluted lung homogenate. Each dilution was assayed in duplicate. The plates were incubated at 37°C, 5% CO_2_ for two hours, with the plates swirled every 15 minutes. Following incubation, the media was aspirated and the Vero E6 cells rinsed twice with sterile PBS. An overlay of DMEM with 2.5% FBS and 1.2% Avicel PH-101 (Sigma Aldrich) was added to each well and the plates were incubated at 37°C, 5% CO_2_. After 2 days, the media was aspirated and the Vero E6 cells were fixed with 10% formaldehyde prior to staining with 0.1% crystal violet. The plaques were counted for each well and viral titers calculated by averaging the PFU/mL calculated for each duplicate according to the equation: PFU/mL = (number of plaques) / (dilution factor × volume diluted lung homogenate added to the well).

### Histology

In these experiments, mice were infected with 1.5 × 10^4^ PFU SARS-CoV-2 MA10 and were euthanized at day 4 post-infection. The lungs were inflated with PBS prior to removal. The right caudal lung lobe was fixed in 10% formalin at room temperature. After 48 hours, the tissue was transferred to 70% ethanol prior to paraffin embedding and staining with H&E by the University of Virginia Research Histology Core. Tissue pathology was scored in a blinded manner according to the American Thoracic Society’s ALI scoring system^23^ which generates a score according to the following criteria: (A) neutrophils in the alveolar space (none (A = 0), 1-5 neutrophils (A = 1), >5 neutrophils (A = 2)), (B) neutrophils in the interstitial space (none (B = 0), 1-5 neutrophils (B = 1), >5 neutrophils (B = 2)), (C) hyaline membranes (none (C = 0)), 1 membrane (C = 1), >1 membrane (C = 2)), (D) proteinaceous debris filling the airspaces (none (D = 0), 1 instance (D = 1), >1 instance (D = 2), and (E) alveolar septal thickening relative to mock-infected control (<2x thickening (E = 0), 2x-4x thickening (E = 1), >4x thickening (E = 2)). The ALI score is calculated as Score = [(20 × A) + (14 × B) + (7 × C) + (7 × D) + (2 × E)]/(number of fields × 100). Three random fields of tissue at 600X total magnification were evaluated for each sample.

### Luminex Analysis

In these experiments, mice were infected with 5.5 × 10^3^ PFU SARS-CoV-2 MA10 and were euthanized at day 4 post-infection. BALF was collected using 0.6 mL of sterile PBS. The BALF samples were centrifuged at 400 ×g for 10 minutes at 4°C. The supernatants were collected and stored at -80°C. Cytokine concentrations in the BALF supernatant were measured using a Luminex 32-plex Mouse panel (MCYTMAG-70K-PX32, MilliporeSigma) that assays Eotaxin/CCL11, G-CSF, GM-CSF, IFN-γ, IL-1α, IL-1β, IL-2, IL-3, IL-4, IL-5, IL-6, IL-7, IL-9, IL-10, IL-12 (p40), IL-12 (p70), IL-13, IL-15, IL-17, IP-10, KC, LIF, LIX, MCP-1, M-CSF, MIG, MIP-1α, MIP-1β, MIP-2, RANTES, TNF-α, and VEGF. The University of Virginia flow cytometry core ran this panel on a Luminex MAGPIX according to manufacturer’s instructions.

### Statistical Analysis

For survival experiments, data were plotted by Kaplan–Meier curve with a log-rank test performed to determine statistical significance between experimental groups. For all other comparisons between experimental groups (evaluating differences in weight loss, viral titers, ALI scoring, and cytokine concentrations) statistical significance determined by non-parametric test. *P<0.05, **P<0.01, ***P<0.001. All statistical analyses were performed using GraphPad Prism software.

## ACKNOWLEDGEMENTS

The authors would like to thank the Petri lab members for their thoughtful discussions of this work particularly Duncan Hart, Nicholas Natale, Dr. Jashim Uddin, and Dr. Barbara Mann. We thank Savannah Brovero and Dr. Mehmet Tanyüksel for technical support. We thank Taylor Harper, University of Virginia Flow Cytometry Core (RRID: SCR_017829), for cytokine measurements. We thank the staff of the University of Virginia Research Histology Core (RRID: SCR_025470), Biorepository and Tissue Research Facility (RRID: SCR_022971), and Center for Comparative Medicine for their technical support.

## AUTHOR CONTRIBUTIONS

C.F. and W.A.P designed experiments with invaluable advice from J.E.A. and H.J.M. C.F. performed experiments and analyzed data. C.F. wrote this manuscript and W.A.P., J.E.A., and H.J.M reviewed and edited the manuscript.

